# Insights into substrate-mediated assembly of the chloroplast TAT receptor complex

**DOI:** 10.1101/2020.05.08.085019

**Authors:** Qianqian Ma, Christopher Paul New, Carole Dabney-Smith

**Affiliations:** Graduate Program in Cell, Molecular, and Structural Biology, Miami University, Oxford, Ohio, USA; Department of Chemistry and Biochemistry, Miami University, Oxford, Ohio, USA

## Abstract

The <u>T</u>win <u>A</u>rginine <u>T</u>ransport (TAT) system translocates fully folded proteins across the thylakoid membrane in the chloroplast (cp) and the cytoplasmic membrane of bacteria. In chloroplasts, cpTAT transport is achieved by three components: Tha4, Hcf106, and cpTatC. Hcf106 and cpTatC function as the substrate recognition/binding complex while Tha4 is thought to play a significant role in forming the translocation pore. Recent studies challenged this idea by suggesting that cpTatC-Hcf106-Tha4 function together in the active translocase. Here, we have mapped the inter-subunit contacts of cpTatC-Hcf106 during the resting state and built a cpTatC-Hcf106 structural model based on our crosslinking data. In addition, we have identified a substrate-mediated reorganization of cpTatC-Hcf106 contact sites during active substrate translocation. The proximity of Tha4 to the cpTatC-Hcf106 complex was also identified. Our data suggest a model for cpTAT function in which the transmembrane helices of Hcf106 and Tha4 may each contact the fifth transmembrane helix of cpTatC while the insertion of the substrate signal peptide may rearrange the cpTatC-Hcf106-Tha4 complex and initiate the translocation event.

**One sentence summary:** Protein subunits of the thylakoidal twin arginine transport complex function together during substrate recognition and translocase assembly.

## Introduction

The <u>T</u>win <u>A</u>rginine <u>T</u>ransport (TAT) system is a unique protein transport pathway found in plant chloroplasts (cpTAT), some plant mitochondria, and plasma membranes of bacteria and archaea (Berks, 2015; Hamsanathan and Musser, 2018; New et al., 2018). In plants, the cpTAT system plays a significant role in transporting several lumen-residing protein photosynthetic machinery subunits (Molik et al., 2001). Null mutations of cpTAT components severely affect chloroplast development having been shown lethal in maize and Arabidopsis seedlings (Das and Martienssen, 1995; Motohashi et al., 2001). In bacteria, TAT is essential for exporting proteins into the periplasm that are involved in energy metabolism, nutrient acquisition, and virulence (Lee et al., 2006; Joshi et al., 2010). The presence of a TAT system in some plant mitochondria may be for the translocation of soluble domains of respiratory membrane proteins (van der Merwe and Dubery, 2007; Carrie et al., 2016).

The most unique feature of TAT is its ability to transport completely folded proteins with a requisite N-terminal twin arginine (RR) motif (Cleon et al., 2015; Cline, 2015) using only the energy derived from the proton motive force (PMF) generated across the lipid bilayer (Cline and Mori, 2001; Bageshwar and Musser, 2007). One question that has yet to be answered since the cpTAT system was first discovered is how folded proteins are transported across a membrane bilayer without ion leakage (Mould et al., 1991; Cline et al., 1992). To address this question and further define the molecular mechanisms of (cp)TAT function, purified chloroplasts and *E. coli* inverted membrane vesicles have been used as model membrane environments. So far, similarities between each system have been identified. In plant and *E. coli* TAT systems, three conserved protein components function together: Tha4, Hcf106, and cpTatC in cpTAT and TatA, TatB, and TatC in *E. coli*. Previous studies suggest that cpTatC (TatC) forms a large stable multi-subunit receptor complex with Hcf106 (TatB) in a strict 1:1 ratio (Bolhuis et al., 2001; Behrendt et al., 2004; Punginelli et al., 2007). Recently, Tha4 was also found constitutively associated with the cpTatC-Hcf106 receptor complex (Aldridge et al., 2014). The size of the receptor complex ranges from 360 and 700 kDa depending on the isolation procedure used with a consensus suggesting approximately six-eight copies of Hcf106 and cpTatC in the receptor complex (Cline and Mori, 2001; Fincher et al., 2003; Oates et al., 2005; Orriss et al., 2007; Behrendt and Bruser, 2014). This hetero-oligomeric complex of cpTatC-Hcf106 functions to recognize and bind the signal sequence of precursor proteins with a RR signal peptide motif (Alami et al., 2003; Gerard and Cline, 2006; Lausberg et al., 2012). After substrate binds with the receptor complex, additional Tha4 is recruited and translocation is initiated (Cline and Mori, 2001). Single particle electron microscopy imaging and biochemical crosslinking experiments have revealed that multiple substrate monomers can bind to the receptor complex (Celedon and Cline, 2012; Berks, 2015). Additional evidence for multiple substrate proteins binding to the receptor complex is that covalently-linked precursors are translocated together (Ma and Cline, 2010, 2013). Despite the wealth of plant and bacterial TAT structural data, a detailed molecular mechanism of this transient protein transport system remains elusive.

Understanding how the receptor complex is organized and determining whether dynamic reorganization occurs during substrate translocation are two pivotal aims of improving mechanistic models of TAT function. Recently, crystal structures of TatC isolated from the thermophilic bacterium *Aquifex aeolicus* (aq) revealed that the six transmembrane domains (TM) of aqTatC adopt an overall glove-shaped structure with a lipid-exposed pocket involved in signal sequence binding (Rollauer et al., 2012; Ramasamy et al., 2013). TatC is the largest protein in the complex with each monomer functioning together as the structural backbone of the translocase. The additional cpTAT proteins, Tha4 and Hcf106, are both membrane proteins with a single TM. NMR-derived structures of bacterial TatA and TatB each have an overall rotated L-shape with an N-terminal TM linked to an amphipathic helix (APH) by a short hinge terminating in an unstructured, soluble C-terminal tail (Hu et al., 2010; Zhang et al., 2014). Recently cpTatC-Tha4 interaction studies found a constitutive interaction between the Tha4 TM and cpTatC TM5 (Aldridge et al., 2014). Additionally, substrate binding has been shown to trigger additional contacts between the Tha4 TM and cpTatC TM4 (in the cavity of the glove-like structure), possibly by forming hydrogen bonds with cpTatC prior to the initiation of transport (Aldridge et al., 2014). However, the role of Hcf106 in complex assembly and how it fits into the Tha4-cpTatC translocase model is unclear.

In this study, our primary goals were to map the contacts between Hcf106 and cpTatC in the receptor complex by disulfide crosslink scanning and to determine whether and how these interactions were affected by precursor signal peptide binding. On the stromal side of the thylakoid membrane, interactions were observed between the N-proximate region of the Hcf106 APH and the N-terminus of cpTatC. Additionally, two stromal loops of cpTatC were shown to crosslink with the Hcf106 APH. In agreement with previous studies in *E. coli*, close contacts between cpTatC TM5 and the Hcf106 TM were also observed in chloroplasts. We identified additional interactions between the Hcf106 TM and cpTatC TM1 and TM2 that are of special interest as some were altered in the presence of precursor. Our work also captured a Tha4-cpTatC-Hcf106 trimer linked by TM domain interactions between each monomer. This observation provides direct evidence for a Tha4-cpTatC-Hcf106 complex during the formation of active translocase. Finally, a model of the cpTAT system is discussed in detail.

## Results

### Hcf106 C-proximate APH forms close contacts with cpTatC stromal domains; Hcf106 N-proximate APH and TM form close contacts with cpTatC TM5

We recently reported a method to study the contribution of Hcf106 in the cpTAT system by integrating *in vitro* translated cys-substituted Hcf106 into isolated thylakoid membranes (Ma et al., 2018). We sought to take advantage of our Hcf106 cys-substitution library to map interactions between Hcf106 and cpTatC in the receptor complex.

We started by investigating the contacts between the C-terminal proximate APH of Hcf106 and the stromal domains of cpTatC as they are most likely to form crosslinks based on the relative positions of the cys residue with respect to the membrane. First, we created single cysteine Hcf106 APH variants (T38C, E48C, R54C, R62C, and I69C) and cpTatCaaa (endogenous cpTatC cysteines replaced with alanine) variants each with a single cysteine substitution in the N-terminal stromal region (S1, residue D78C) or one of the two stromal loops (S2 and S3, residues T164C and Q244C, respectively) (Fig. 1A). The region of Hcf106 spanning from the N-terminus to N34 was predicted to partition into the thylakoid membrane (New et al., 2018), so we substituted cys for APH residues from T38 and beyond in our study. [^3^H]cpTatCaaa D78C (S1), -T164C (S2), and -Q244C (S3) were separately imported into chloroplasts and secreted into the thylakoid membrane by cpSec. We then integrated the unlabeled Hcf106 single cys-variants into cpTatCaaa cys variant containing thylakoids and promoted disulfide crosslink oxidation with copper 1,10-phenanthroline (CuPhen; see Materials and Methods). The negative controls lacked integrated Hcf106 cys variants. cpTatC (29 kDa)-Hcf106 (28 kDa) crosslinks appeared as a 57 kDa adduct (Fig. 1B-D). Crosslink formation between the Hcf106 APH and the N-terminal (D78C) or stromal loop 2 (T164C) regions of cpTatC increased as the Hcf106 APH cys substitutions were placed closer to the C-terminus (Fig. 1B). cpTatC stromal loop S3 (Q244C)-Hcf106 APH interactions were the most prominent for E48C and R54C (Fig. 1B, lanes 15-16). In summary, close contacts are formed between the Hcf106 C-proximate APH region and cpTatC S1 and S2 at rest.

**Figure 1.**
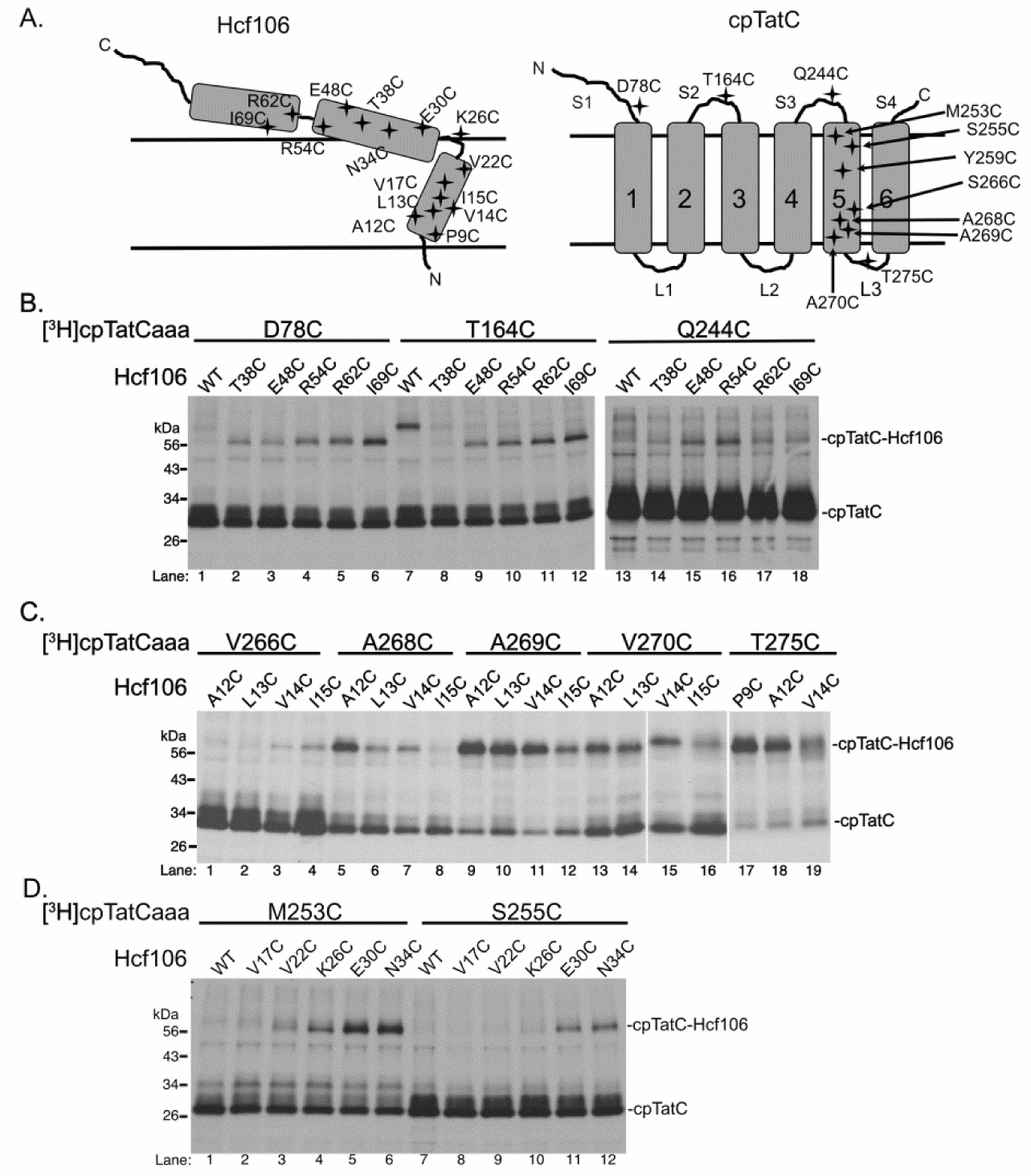
Hcf106 C-proximate APH forms close contacts with cpTatC stromal domain; Hcf106 N-proximate APH and TM form contacts with cpTatC TM5. Thylakoid membranes containing single cys variants of [^3^H]cpTatCaaa were incubated with unlabeled Hcf106 single cys variants and subjected to oxidative crosslinking (see Materials and Methods). Samples were analyzed by non-reducing SDS-PAGE and fluorography. **A** Topology models of Hcf106 and cpTatC. **B** Interaction between Hcf106 C-proximate APH and cpTatC with cys residues in the stromal loops (S1, S2, and S3). **C** Interactions between Hcf106 central transmembrane domain and cpTatC TM5. **D** Crosslinking between Hcf106 hinge region and cpTatC TM5.

We also determined the interactions between cpTatC and the Hcf106 TM region. In *E. coli*, the TatB TM was repeatedly shown to form contacts with TatC TM5 (Kneuper et al., 2012; Rollauer et al., 2012; Alcock et al., 2016). However, the relative positions of these two TM helices are unclear. So, we need to precisely determine their relative positions to better define how the translocase is assembled. We employed our cysteine scanning method to detect contacts between cpTatC TM5 (V266C, A268C, A269C, V270C) and the Hcf106 TM (A12C, L13C, V14C, I15C) (Fig. 1C). cpTatC V266C (Fig. 1C, lanes 1-4) showed minimal interaction with each Hcf106 TM variant. cpTatCaaa A268C (sulfhydryl side chain predicted to face TM6) showed a relatively strong interaction with Hcf106 A12C and mild contacts with L13C and V14C (Fig. 1C, lanes 5-7). cpTatCaaa A269C and V270C formed the strong interactions with the Hcf106 TM indicates that these residues orient towards the Hcf106 TM (Fig. 1C, lanes 9-16). Finally, interaction between cpTatC lumen loop 3 (T275C) and the Hcf106 TM was examined. The Hcf106 TM cys variant A9C formed the strongest contacts with cpTatCaaa T275C which suggests a prominent interaction between cpTatC lumen loop 3 and Hcf106 N-terminal region in the receptor complex (Fig. 1C, lanes 17-19).

Finally, we mapped interactions between the stroma proximate portion of cpTatC TM5 (M253C and S255C) and Hcf106 C-proximate TM (V17C, V22C), hinge (K26C), or N-proximate APH (E30C, N34C) regions (Fig. 1D). Both cpTatCaaa M253C and S255C interacted with Hcf106 E30C and N34C (Fig. 1D, lanes 5, 6 and 11, 12), which suggests that the N-proximate APH of Hcf106 is partially inserted into the membrane in a similar manner to the Tha4 APH (Aldridge et al., 2012).

Taken together, the disulfide mapping presented here indicates that the Hcf106 C-proximate APH forms close contacts with the cpTatC stromal domains. Additionally, the Hcf106 TM seems to rest parallel with cpTatC TM5 based on the contacts formed between Hcf106 and the short lumen loop region 3 that links cpTatC TM5 to TM6.

### The cpTatC-Hcf106 adduct appears as a 57 kDa complex

To verify that the 57 kDa crosslink product represents disulfide bond formation between cpTatC and Hcf106, we used anti-FLAG immunoprecipitation to capture radiolabeled or unlabeled Hcf106 A12C-FLAG that was crosslinked with [^3^H]cpTatCaaa A269C. Unlabeled, wild type (lacks endogenous cysteines) Hcf106-FLAG did not capture any [^3^H]cpTatCaaa A269C (Fig. 2, lanes 1-2). Hcf106 A12C-FLAG immunoprecipitated with [^3^H]cpTatCaaa A269C in a ~57 kDa adduct (Fig. 2, lanes 3-4) that had a greater observed intensity when both cpTatC and Hcf106 were radiolabeled (Fig. 2, lanes 5-6). When we used a shorter exposure time, we clearly differentiated the 57 kDa cpTatC-Hcf106 adduct from the 56 kDa Hcf106-Hcf106 band (Fig. 2, lane 10). Thus, co-purification of radiolabeled cpTatCaaa with unlabeled Hcf106-FLAG confirmed that the 57 kDa complex is an adduct of Hcf106 and cpTatC.

**Figure 2.**
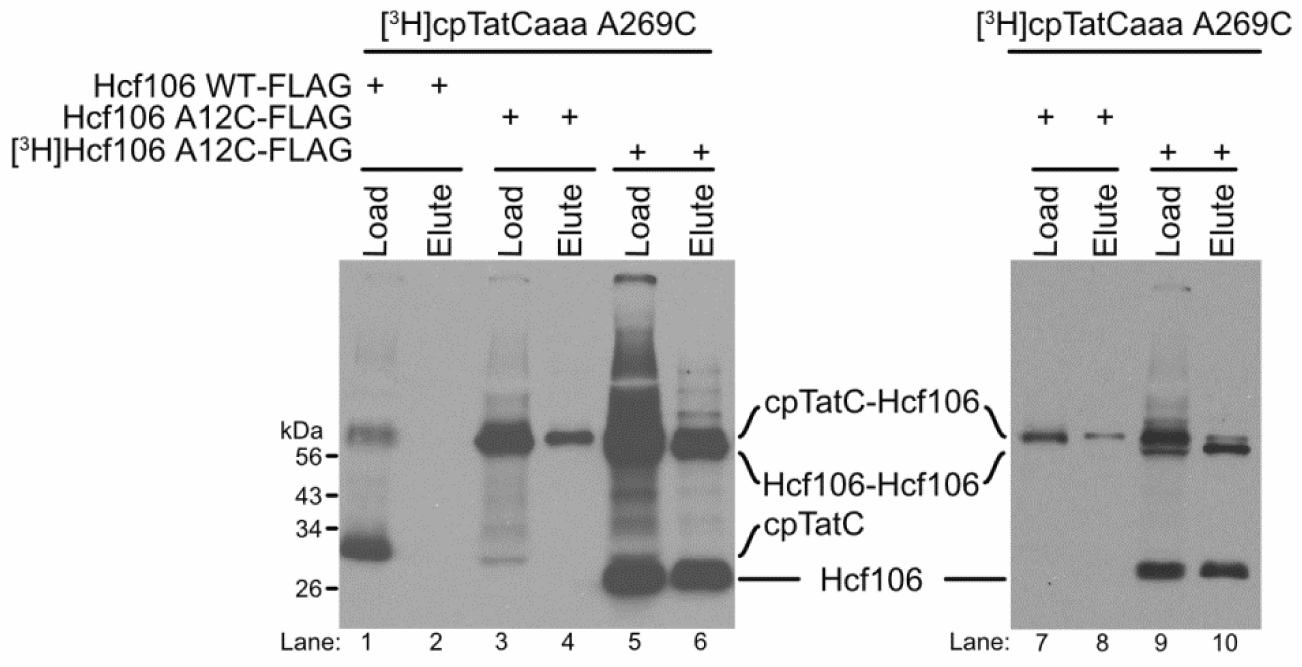
The 57 kDa band is an adduct of cpTatC-Hcf106. [^3^H]cpTatCaaa A269C was imported into thylakoids followed by integration of radiolabeled or unlabeled Hcf106 variants as indicated. After oxidative crosslinking, samples were subjected to immunoprecipitation with beads labeled with antibodies against the FLAG tag. Lanes 7-10 are diluted samples from lanes 3-6. Samples were analyzed by SDS-PAGE and fluorography under non-reducing conditions (see Materials and Methods).

### Hcf106 TM contacts both cpTatC TM5 and Tha4

Previously, a Tha4 cys substitution (F3C) located in N-terminal TM region was shown to form a constitutive interaction with cpTatC TM5 V270C (Aldridge et al., 2014). In this study, we identified similar interactions between Hcf106 TM positions A12C, L13C, V14C, and I15C with cpTatC TM5 A269C (Fig. 1C, lanes 9-12). As such, we sought to capture all three cpTAT components in one complex after integration into thylakoid membranes. Tha4 F3C and Hcf106 G6C interacted weakly with [^3^H]cpTatCaaa A269C (Fig. 3A, lanes 2-3) while Hcf106 V14C formed the most prominent interaction with this cpTatC residue (Fig. 3A, lane 4). So, we generated double cys substituted Hcf106 G6C V14C to capture the Tha4 with one cys and cpTatC with the other. Hcf106 G6C V14C formed a complex with [^3^H]cpTatCaaa A269C equivalent to a ~90 kDa cpTatC-Hcf106-Hcf106 trimer (Fig. 3A, lane 5). When Tha4 F3C was added to the reaction, we detected a ~68 kDa band (Fig. 3A, lane 6) that corresponds to cpTatC-Hcf106-Tha4 and disappears after reduction of the disulfide bonds (Fig. 3A, lane 13). To verify that this ~68 kDa band is a cpTatC-Hcf106-Tha4 adduct, we used anti-FLAG immunoprecipitation to capture Hcf106 G6C V14C-FLAG crosslinked with [^3^H]cpTatCaaa A269C and Tha4 F3C. Hcf106 G6C V14C-FLAG coprecipitated with [^3^H]cpTatCaaa and appeared as a ladder of cpTatC-Hcf106 adducts during SDS-PAGE analysis (Fig. 3B, lane 2). When Tha4F3C was present, a ~68 kDa band was present after co-IP (Fig. 3B, lane 4) that suggests we captured a complex of the three cpTAT components.

**Figure 3.**
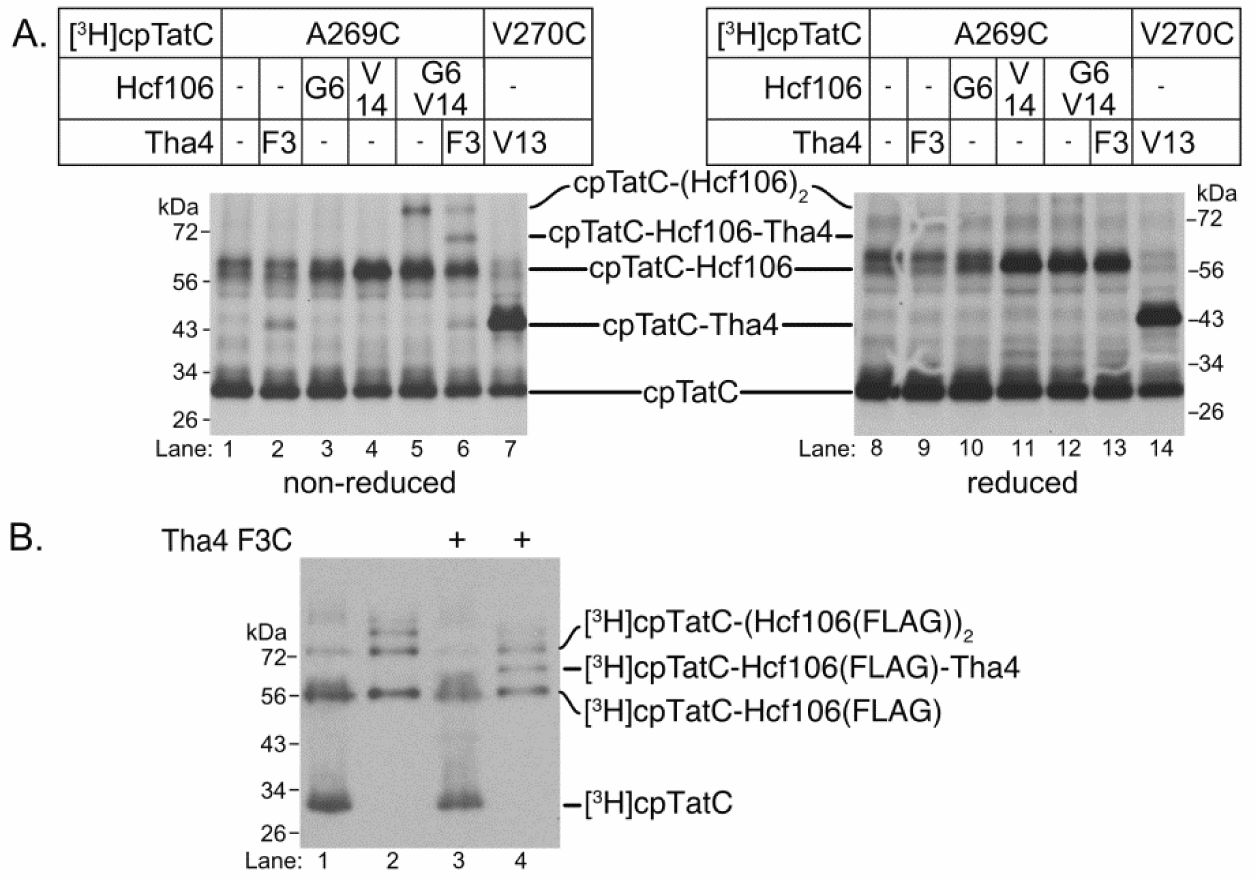
Recombinant cpTatC, Hcf106, and Tha4 exist in one complex and can be crosslinked through a dual cys substituted Hcf106. **A** Thylakoid membranes containing [^3^H]cpTatCaaa single cys variants (A269C, V270C) were incubated with unlabeled Hcf106 (G6C, V14C, or G6C V14C) or Tha4 (F3C, V13C). Following the addition of Hcf106 and/or Tha4, crosslinking was induced by CuPhen (see Materials and Methods). Samples were analyzed by non-reducing and reducing SDS-PAGE and fluorography. Predicted dimers and trimers are indicated. **B** Immunoprecipitation of Hcf106(FLAG)-cpTatC-Tha4 trimer. Unlabeled cpTatCaaa A269C was imported into thylakoids followed by the integration of radiolabeled Hcf106 G6C V14C-FLAG (lanes 1-4) in the absence (lanes 1-2) or presence (lanes 3-4) of Tha4 F3C followed by crosslinking. Non-reduced samples (lanes 1 and 3) were immunoprecipitated with αFLAG-beads (lanes 2 and 4) and analyzed by SDS-PAGE and fluorography.

### Hcf106 TM also contacts cpTatC TM1 and TM2

Our initial crosslinking data of the cpTatC-Hcf106 depicted their organization during the resting state (Fig. 1). However, substrate translocation is a rapid and dynamic process and we asked whether there could be subtle movement of Hcf106 within the cpTatC cavity. We used a series of cpTatCaaa variants with single cysteine substitutions in TM1 (I98C), TM2 (L144C, L145C), and TM4 (S228C) together with a control in TM5 (A269C) (Fig. 4A) to investigate if Hcf106 shifts changes from an at rest (Fig. 1) translocase active position. These residue locations were chosen based on the structure of bacterial TatC in which TM1/2/4/5 form the membrane-facing pocket and TM3/6 maintain the back of the glove-like structure (Fig. 4A) (Ramasamy et al., 2013). Prior to oxidative crosslinking, *in vitro* translated unlabeled precursor (V-20F-18C)tOE17 was added to cpTatCaaa and Hcf106 cys variant integrated thylakoids under moderate illumination to generate the active form of the translocase. Of the cpTatC cys variants tested, A269C in TM5 most readily crosslinked with the Hcf106 TM (Fig. 1C). We identified additional contacts between Hcf106 TM and cpTatC TM1 and TM2. cpTatCaaa I98C (TM1) and L144C (TM2) each crosslinked with Hcf106 A12C in the absence of precursor (Fig. 4B, lanes 1, 3) and formation of these interactions decreased when precursor was present (Fig. 4B, lanes 2, 4).

**Figure 4.**
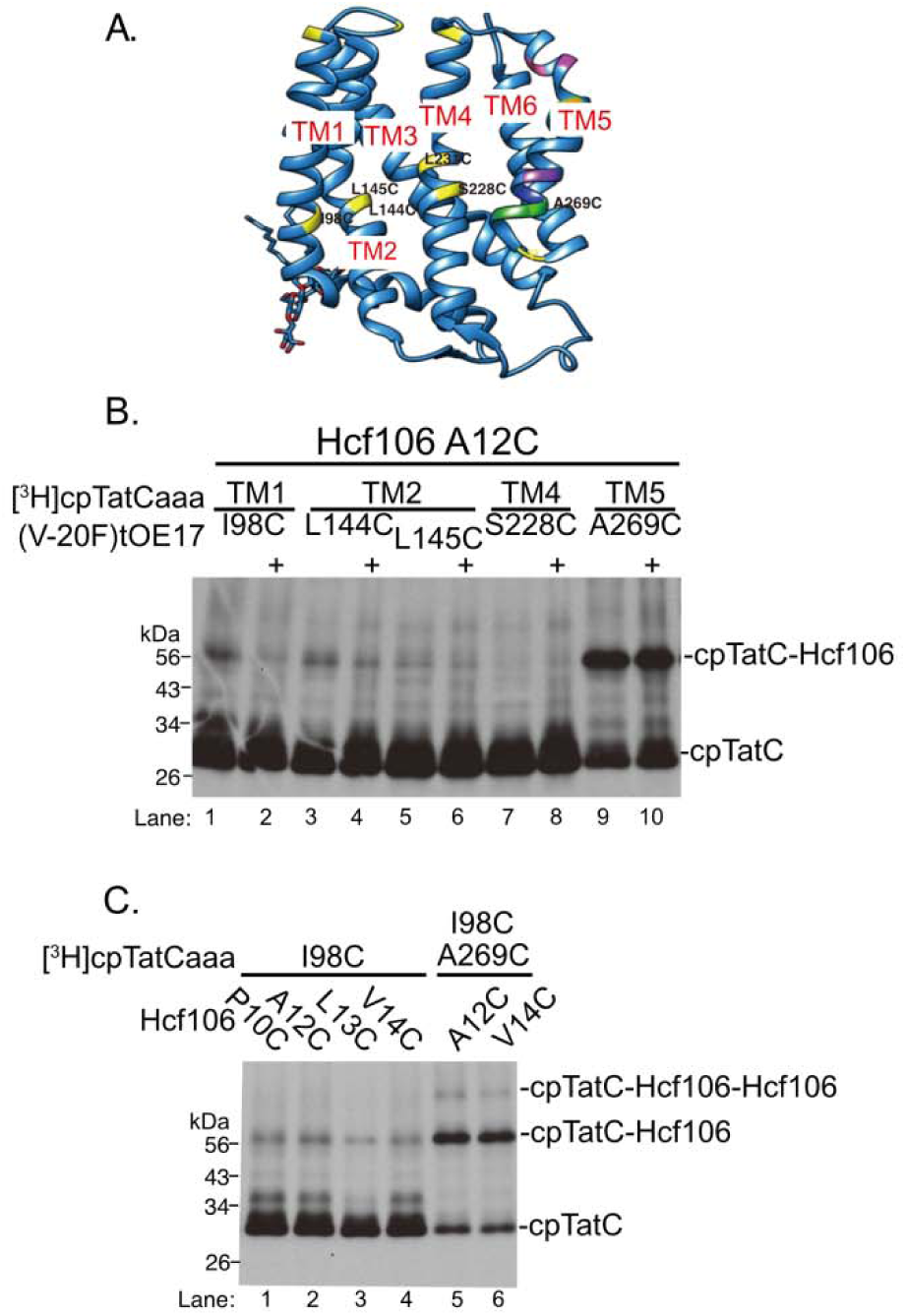
The transmembrane domain of Hcf106 forms close contacts with cpTatC TM1, TM2, and TM5. [^3^H]cpTatCaaa cys variants were imported into isolated chloroplasts and crosslinked with unlabeled Hcf106 integrated thylakoids. Crosslinking was conducted in the absence and presence of precursor (V-20F-18C)tOE17. Samples were analyzed by non-reducing SDS-PAGE and fluorography (see Materials and Methods). **A** Cartoon model of cpTatC from garden pea based on homology with *A. aeolicus* TatC crystal structure (Rollauer et al., 2012) labeled with cys substitutions in yellow (see Materials and Methods). **B** Interactions between Hcf106 A12C and [^3^H]cpTatCaaa TM1, TM2, TM4, and TM5 in the absence or presence of precursor, (V-20F)tOE17. **C** Contacts between Hcf106 A12C and cpTatCaaa TM1 I98C and TM5 A269C in the absence of precursor. Double cys variant cpTatCaaa I98C A269C forms a trimer of cpTatC-Hcf106-Hcf106.

Hcf106 A12C interactions the other cpTatCaaa variants were unchanged regardless of precursor (Fig. 4B, lanes 3-10). To confirm the newly identified Hcf106 TM-cpTatC TM1 interaction additional Hcf106 TM cysteine variants were tested in the absence of precursor. Hcf106 P10C, L13C, and V14C consistently crosslinked with cpTatC I98C in the absence of precursor (Fig. 4C, lanes 1-4). This data confirms a constitutive interaction between the Hcf106 TM and cpTatC TM1. cpTatC TM1 and TM5 each formed crosslinking products with Hcf106. In terms of receptor complex organization and the overall 1:1 ratio of cpTatC and Hcf106, these results indicate that each cpTatC either contacts a single Hcf106 via TM1 and TM5 or that one cpTatC could be in close contact with two Hcf106 monomers at the same time. A cpTatC double cysteine mutant was constructed to test the latter scenario. When crosslinking cpTatC I98C A269C to Hcf106 A12C or V14C, we observed additional bands that show the ability of cpTatC to contact two Hcf106 monomers at the same time (Fig. 4C, lanes 5-6). Taken together, these data indicate the presence of two Hcf106 monomers near one cpTatC monomer in the absence and presence of precursor.

### The interaction between cpTatC TM5 and Hcf106 TM is enhanced during precursor transport

Disulfide crosslinking between cpTatCaaa I98C (TM1) and Hcf106 A12C was substrate dependent; however, the interaction between cpTatCaaa A269C and Hcf106 A12C was seemingly unchanged by precursor. Thus, we repeated the precursor addition experiment with [^3^H]cpTatCaaa V270C where the sulfhydryl is closer to the concave face of cpTatC. To further maximize differences in crosslink formation, we used a lower CuPhen concentration [0.5 mM] and a shorter oxidation period (2 min). Prior to the addition of precursor (0 min), a faint crosslink product was observed (Fig. 5A, lane 3). After 5 min, a pronounced increase in crosslinked product was observed (Fig. 5A, lane 4) which decreased after 15 min (Fig. 5, lane 5). Successful transport without crosslinking was confirmed by thermolysin degradation of precursors that remained outside of the thylakoid lumen (Fig. 5, lanes 1-2).

**Figure 5.**
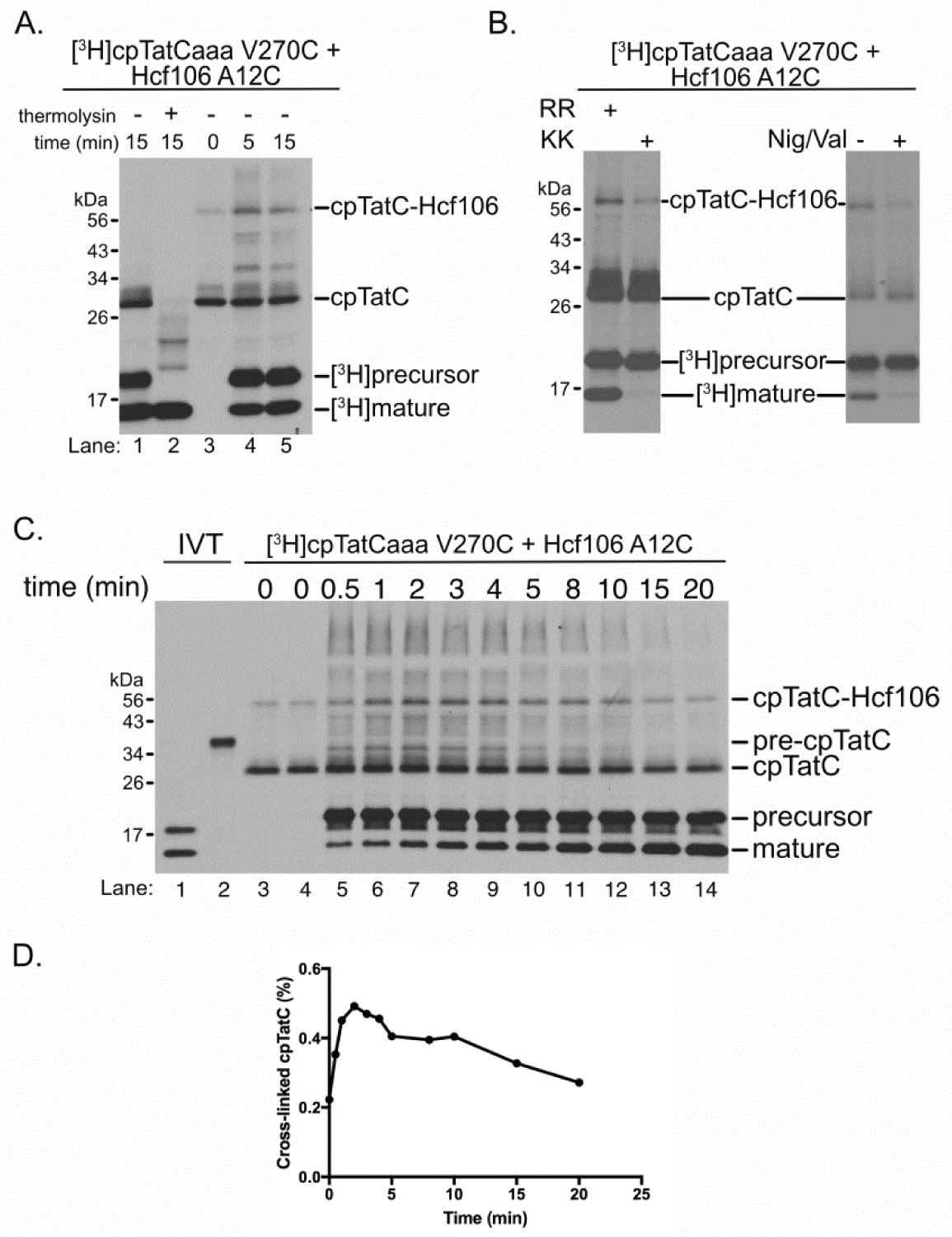
Interaction between cpTatC TM5 and Hcf106 TM A12C was enhanced during precursor transport. Thylakoid membranes containing [^3^H]cpTatCaaa V270C were incubated with unlabeled Hcf106 A12C followed by addition of precursor [^3^H](V-20F)tOE17. Transport was induced by illumination at 25 °C. Samples were crosslinked with 0.5 mM CuPhen for 2 min at different time points after start of transport as indicated. Transport was then quenched with 3-fold volume of 50 mM NEM. **A** Lanes 1 and 2, substrate was transported for 15 min and subjected to thermolysin treatment to check for complete transport. Lanes 3-5, samples were taken before transport (0 min) and 5 and 15 min after transport. **B** [^3^H]cpTatCaaa V270C interaction with Hcf106 A12C is decreased in the presence of KKtOE17 and nigericin/valinomycin (Nig/Val). **C** IVT corresponds to *in vitro* translated precursor and cpTatC variant. Crosslinked samples were taken from 0.5 to 20 min after the start of transport and subjected to crosslinking (see Materials and Methods). **D** cpTatC dimer/total cpTatC were quantified with ImageJ software and plotted to show % crosslinked cpTatC over unit time.

To confirm that this precursor-dependent cpTatC-Hcf106 interaction is “on pathway”, we repeated the experiment in the presence of transport incompetent precursor KKtOE17 (RR motif replaced with twin lysines) and in the absence of PMF after dissipation by nigericin and valinomycin (Fig. 4B). The interaction of cpTatC-Hcf106 was not promoted in the presence of non-transportable precursor and dissipation of the PMF inhibited transport and diminished the formation of cpTatC-Hcf106 interactions (Fig. 6B). These observations indicate that interaction between cpTatC TM5 and Hcf106 TM occurs during cpTAT assembly.

**Figure 6.**
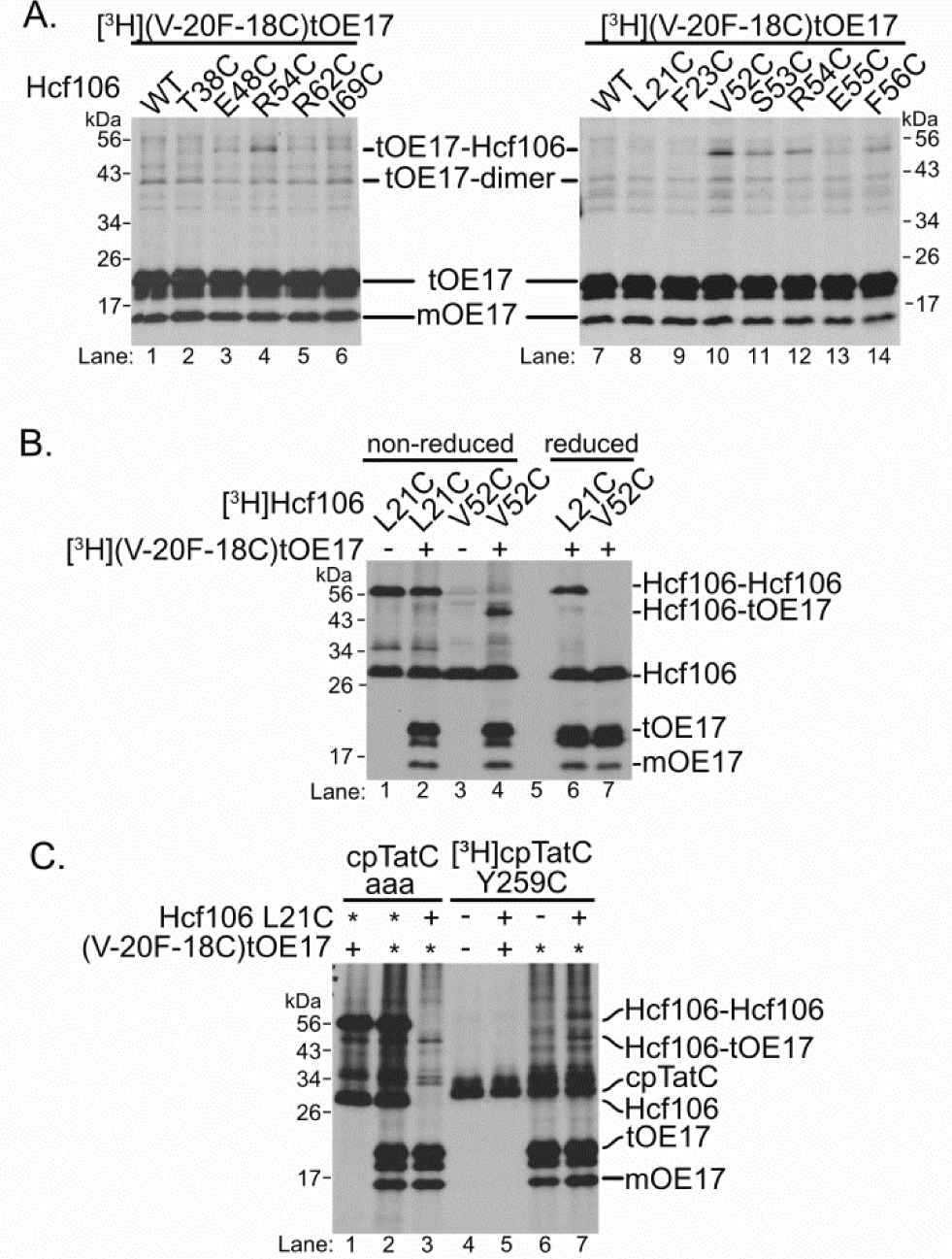
Precursor signal peptide interacts with both Hcf106 APH and TM forming a ~45 kDa adduct. **A** Substrate protein signal peptide interacts with Hcf106 APH. Thylakoid membranes containing unlabeled Hcf106 cys variants were incubated in the presence [^3^H](V-20F-18C)tOE17 for 2 min followed by crosslinking (see Materials and Methods). Samples were analyzed by SDS-PAGE and fluorography under non-reducing conditions. **B** [^3^H]Hcf106 L21C and V52C interacted with [^3^H](V-20F-18C)tOE17 through oxidized crosslinks (lanes2, 4) that were depleted by reducing sample buffer (lanes 6-7). **C** Substrate protein signal peptides interact with Hcf106 L21C. Unlabeled cpTatCaaa or [^3^H]cpTatCaaa Y259C were imported into chloroplasts to help incorporated additional Hcf106 into receptor complexes. Labeled (*) or unlabeled (+) Hcf106 L21C or V52C variant integrated thylakoids received labeled (*) or unlabeled (+) (V-20F-18C)tOE17 to capture precursor-Hcf106 crosslink formation. Thylakoid samples lacking Hcf106 and/or precursor are indicated with (-).

Finally, to correlate precursor transport with receptor complex interactions, we conducted a detailed time course experiment and plotted the percentage of crosslinked cpTatC over time (Fig. 5C-D). We observed an increase of crosslinked Hcf106-cpTatC as the reaction time progressed for ~4 min (Fig. 5C, lanes 3-9). During the 5-10 min range, cpTatC-Hcf106 dimer began to decrease (Fig. 5C, lanes 10-12) and was near pre-transport levels after 15 min (Fig. 5C, lanes 13 and 14). Our time course data agrees with Tha4 transport kinetics that showed maximal transport at ~3 min (Fig. 5D) (Celedon and Cline, 2012). Together, these results show that the interaction between cpTatC TM5 and the Hcf106 TM is highly dependent on precursor transport.

### Both the Hcf106 C-proximate APH and TM interact with precursor near RR region of the signal peptide

Precursor signal peptide binding to the receptor complex initiates the assembly of the TAT translocase (Blaudeck et al., 2001; Cline and Mori, 2001; Alami et al., 2003; Ma and Cline, 2013) likely by interaction with cpTatC S1 and S2 domains (Ma and Cline, 2013). As shown above, S1 and S2 form close contacts with the Hcf106 APH so we expected the signal peptide RR region to also interact with the Hcf106 APH. To investigate this, we used the precursor protein (V-20F-18C)tOE17 with a cys substitution near the RR motif (Fig. 6A). Among the Hcf106 single cys-variants T38C-I69C, R54C was formed the most interactions with (V-20F-18C)tOE17 (Fig. 6A, lane 4). We then narrowed down the location of interaction with Hcf106 by testing cys substitutions in V52-F56 and found that V52C formed the strongest interaction with precursor (Fig. 6A, lane 10).

We also explored whether Hcf106 L21C (stroma proximate TM) interacted with signal peptide. Initially, we did not see any significant interaction between L21C and the precursor signal peptide (Fig. 6B, lanes 1-2). However, we previously demonstrated that integrated Hcf106 competes with endogenous Hcf106 for assembly into the receptor complex (Ma et al., 2018). To improve the interaction efficiency, unlabeled cpTatCaaa was imported into chloroplasts to increase incorporation of exogenous [^3^H]Hcf106 into receptor complexes. During transport in the presence of [^3^H]Hcf106 L21C or [^3^H](V-20F-18C)tOE17, we detected a ~48 kDa that is equivalent to Hcf106-precursor (Fig. 6C, lanes 1, 3) and the band intensity was enhanced if both Hcf106 and precursor were radiolabeled (Fig. 6C, lane 2). We also detected this adduct if we imported [^3^H]cpTatC Y259C but only when Hcf106 L21C was also present (Fig. 6C, lanes 6-7).

Together, these results suggest a progression for precursor signal peptide binding. First, the precursor signal peptide forms contacts with the Hcf106 APH and cpTatC stromal proximate domains. The signal peptide is then inserted into the concave face of the cpTatC-Hcf106 complex where it can interact with the Hcf106 TM and cpTatC (Gerard and Cline, 2007). This movement of the signal peptide may play a significant role in inducing a conformational change in the receptor complex that ultimately assists the translocation of the precursor mature domain.

## Discussion

In this study, we sought to map Hcf106-cpTatC interactions in the receptor complex at rest and how these contacts change in the presence of precursor signal peptide and translocase assembly. Ultimately, we wanted to better define Hcf106 function in cpTAT beyond precursor binding (Gerard and Cline, 2007). Previous work showed that Tha4 formed close contacts with cpTatC TM5 in non-stimulated membranes (Aldridge et al., 2014). When precursor and PMF was present, the Tha4 TM formed crosslinks with cpTatC TM4 (Aldridge et al., 2014) which led the authors to propose a model of cpTAT where Tha4 enters and possibly forms the translocation element in the enclosed center of precursor bound receptor complexes (Aldridge et al., 2014). However, this model lacked a defined role for Hcf106 during translocase assembly. We showed that Hcf106 TM interactions with cpTatC TM1/2/5 change in the presence of precursor (Fig. 1, 4) like TatB-cpTatC in *E. coli* TAT (Kneuper et al., 2012; Alcock et al., 2016). These interactions suggest a face-to-face dimeric cpTatC model (Fig. 7B) that is supported by prior work where S1 domains of separate cpTatC monomers simultaneously crosslinked a linked precursor pair (Aldridge et al., 2014). This model assumes a dimer of trimeric receptor complexes where Hcf106/Tha4 function as a gate located between cpTatC TM1 and TM5 (Aldridge et al., 2014). Once receptor complex binds with precursor, the Hcf106 TM switches interaction preference from cpTatC TM1 to TM5 (Fig. 5). Previously, a similar model of *E. coli* TAT was proposed in which precursor signal peptide binding decreased TatB TM affinity for TatC TM5/6 domain leading to TatA occupation of this binding site (Alcock et al., 2016). Our data differs from the *E. coli* model as cpTAT precursor binding depleted the cpTatC TM1-Hcf106 TM interaction, not cpTatC TM5-Hcf106 TM interactions. Although both cpTAT and *E. coli* TAT are quite similar, this may be an emerging functional difference between these two systems.

**Figure 7.**
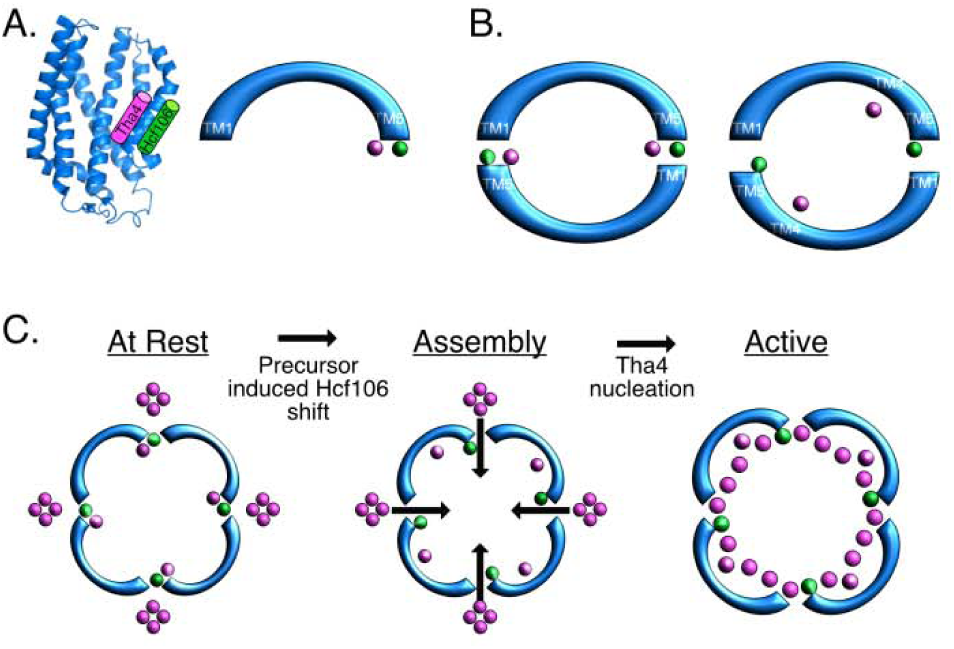
Model of Hcf106-Tha4-cpTatC complex at rest and during translocase assembly. **A** Side view (left) and top-down view (right) of cpTatC (blue) modeled with Hcf106 (green) and Tha4 (pink). The TMs of Hcf106 and Tha4 compete to bind in the same position near cpTatC TM5. **B** Dimeric cpTatC-Hcf106-Tha4 at rest (left) and during translocase assembly (right). In the resting state, Hcf106 remains close to cpTatC TM5 as well as TM1 and TM2 of an adjacent cpTatC. During translocation, Tha4 moves to the interior of the cpTAT cavity and Hcf106 affinity for adjacent cpTatC TM1 decreases. **C** Model of cpTAT assembly based on a tetramer of receptor complex trimers. Binding of precursor signal peptide diminishes Hcf106 interactions with adjacent cpTatC TM1 and promoting Tha4 binding with cpTatC TM4. Additional Tha4 from the thylakoid membrane enters through the open “gates” into the center of the cpTAT annulus. Finally, Tha4 organizes in the central chamber possibly by nucleation with the Tha4 bound to cpTatC TM4 and Hcf106.

At rest Hcf106-cpTatC interactions are changed upon addition of precursor, so we also wanted to better define the points of contact between signal peptide and receptor complex component proteins. Signal peptide interactions with cpTatC and Hcf106 have been previously identified (Gerard and Cline, 2006; Ma and Cline, 2013). S1 and S2 are functionally important regions of cpTatC and have been implicated in precursor signal peptide binding (Holzapfel et al., 2007; Zoufaly et al., 2012; Ma and Cline, 2013). Another binding event was captured between cpTatC TM5 V270C and a signal peptide cysteine substitution 7 residues upstream of the mature domain (−7) (Aldridge et al., 2014). To expand on Aldridge et al. (2014) we substituted a cysteine at position −18 in the hydrophobic (H) domain of signal peptide and observed crosslink formation with cpTatC TM5 (Fig. 6C) and, surprisingly, both Hcf106 TM (Fig. 6B, 6C) and APH (Fig. 6A) domains. These results are interesting because Hcf106 APH (V52C) and TM (L21C) are approximately 30 Å apart so simultaneous contacts at the same location on the signal peptide H-domain highly unlikely. Two possible scenarios explain this binary interaction. In the first scenario, precursor signal peptide undergoes two stages of binding where Hcf106 C-proximate APH functions as a primary point of contact and H-domain hydrophobicity drives deeper insertion in proximity to cpTatC TM5 and Hcf106 TM. This agrees with prior work that showed an exposed receptor complex precursor binding site that initially promotes shallow signal peptide insertion by RR-dependent electrostatics that are surpassed by the hydrophobic affinity the H-domain resulting in deeper signal peptide insertion near the Hcf106 N-proximate APH. If that is the case, we should expect to see that precursor crosslinking with the Hcf106 APH occurs earlier than with Hcf106 L21C, but this remains to be shown. The second interpretation of the binary signal peptide interaction with Hcf106 is that the Hcf106 TM (L21C) is brought closer to the APH (V52C) by a transport induced conformational shift. However, this is unlikely as shifting the hydrophobic Hcf106 TM to the stromal face of the thylakoid membrane is energetically unfavorable.

It is still not fully understood how the cpTAT translocase initiates transport after binding folded precursor, but all models propose that Tha4 plays a significant role in this process. Tha4 is generally considered to be the core translocation conducting element by organizing into a pore-like structure that has direct contact with the substrate protein during the translocation event (Dabney-Smith et al., 2006; Pal et al., 2013). Tha4 has also been shown to undergo a conformational shift that is believed to disrupt the membrane bilayer through hydrophobic mismatch between the oligomer of short Tha4 TMs and the thylakoid lipids (Aldridge et al., 2012). However, Tha4 serving as the only component of the translocation pore is unlikely because molecular dynamics (MD) simulations suggest that oligomers of TatA alone are highly unstable in a lipid bilayer (Rodriguez et al., 2013). Based on the similar tilted structures of Tha4 and Hcf106 as well as their comparable sites of contact mapped onto cpTatC (Fig. 7), it is highly possible that Hcf106 acts as a nucleation center to recruit Tha4 to the center of the receptor complex (Fig. 7). Further support of this model is evidence that the mature, folded domain of the precursor interacts with TatB in *E. coli* (Maurer et al., 2010). While we were writing this manuscript, a novel function of Hcf106 was identified. Ouyang et al., found Hcf106 reserve phase separation and facilities substrate translocation by interacting with kyrin-repeat proteins, STT1, STT2 via the loosely structured C tail (Ouyang et al., 2020). This finding further highlights the important role of Hcf106 in cpTAT translocation and distinguishes chloroplast TAT from bacterial TAT system.

Based on our Hcf106-Hcf106 and Hcf106-Tha4 interaction data and the models of Aldridge et al. (2014) and Alcock et al. (2016), we propose that the TMs of Hcf106 and Tha4 have a sustained competition for binding to cpTatC TM5/6 (Fig. 7). At rest, Hcf106 and Tha4 TMs in one trimer link with cpTatC of the adjacent trimer forming a protein annulus that maintains the membrane barrier in the center of the complex (Fig. 7B-C). The resting state bacterial TAT model by Alcock et al. (2016) also favors a large oligomeric TatBC ring complex where TatB functions as a link between each TatC monomer (Alcock et al., 2016). The deep insertion of the signal peptide H-domain into the receptor complex pocket causes a conformational change that shifts Hcf106 such that its interactions with cpTatC TM1 are diminished (Fig. 7B-C). This opens the “gate” for Tha4 influx into center of precursor bound receptor complex (Fig. 7C). The model of bacterial TAT transport from Alcock et al. (2016) proposed a similar mechanism in which TatA replaces TatB at TatC TM5/6. Recent work has also shown that TatB undergoes a conformational change upon signal peptide binding that led to TatA oligomer formation (Huang et al., 2017). Our model of cpTAT also employs the PMF as a means of neutralizing the charge of the conserved glutamate (E10) in Tha4 and E11 in Hcf106. Thus, the generation of a PMF promotes packing of the N-termini of multiple Tha4 TM forming a hydrophilic region at the lumen face of the translocase that provides a “pulling force” for the precursor protein to be transported. The APH regions of Tha4 and Hcf106 then undergo closer interactions with each other due to the tighter packing of the TMs which then brings the precursor to the vicinity of the translocation center. Ultimately, the precursor protein would be translocated in the already weakened-membrane region confined to the interior of the receptor complex. Future studies using quantitative real-time methods to capture the dynamic and transient moment of precursor protein binding to the receptor complex in native membrane are needed to provide more insight into the mechanism of the cpTAT pathway.

The physiological significance of the observed higher ordered cpTAT receptor complex is still unclear. Specifically, further clarification is needed to show whether the functional cpTAT receptor complex is an aggregate of individual cpTatC-Hcf106-Tha4 complexes that each function independently or that higher ordered receptor complexes are an evolutionary solution of cpTAT to improve net translocation efficiency. We favor the latter scenario and think the actual translocation site is in the center of a structure which consists of at least a dimer of cpTatC-Hcf106-Tha4 trimers (Fig. 7B) while a trimer or tetramer of trimers is more probable (Fig. 7C). Our tetrameric model of cpTatC-Hcf106-Tha4 receptor complexes is like the molecular model of bacterial TatABC organization presented by Alcock et al. (2016). There are at least two benefits to this arrangement. First, it confines thylakoid membrane distortion to a small area encircled by the backbone protein cpTatC thus preventing ion leakage and catastrophic membrane disruption Second, Tha4 could be simultaneously recruited by each cpTatC-Hcf106-Tha4 trimer which would promote rapid assembly of the translocase and equal opportunities of precursor transport at each receptor complex unit (Fig. 7C).With the rapid development of cryo electron tomography and cryo CLEM (Correlative Light and Electron Microscopy) techniques, we would like to see a real picture of how TAT behaves in a native thylakoid membrane. trg

## Materials and Methods

### Generation of cysteine-substituted cpTatC, Hcf106, Tha4, and substrate proteins

cpTatC single-cysteine constructs (D78C, I98C, T164C, Q244C, V270C, T275C in the cys-less background, cpTatCaaa) and additional cpTatC single-cysteine variants were made by QuikChange site-directed mutagenesis (Agilent Technologies) (Ma and Cline, 2013; Aldridge et al., 2014; Ma et al., 2018). These constructs were prepared from a template containing the transit peptide and first 13 residues of the mature domain of the pea small subunit of ribulose-1,5-bisphosphate carboxylase/oxygenase (GenBank accession no. AAA33685) that was fused to the N-terminus of the complete mature precursor sequence for pea (*P. sativum*) cpTatC which was cloned into the pGem4Z vector. The three native cysteines of cpTatC were mutated to alanine. Natively lacking cysteine, both Tha4 proteins (Tha4 XnC) and Hcf106 proteins (Hcf106 XnC) with cysteine substitutions were generated previously by the same method (Dabney-Smith et al., 2003; Aldridge et al., 2012; Ma et al., 2018). The (V-20F)tOE17 precursor protein, which is a modified form of the OE17 precursor protein from maize containing a truncated signal peptide with a valine to phenylalanine substitution, was used to generate single cysteine substitutions and was a gift from Dr. Ken Cline (University of Florida).

### Preparation of chloroplasts and thylakoids

Intact chloroplasts were prepared from 10-12-day old pea seedlings (*Pisum sativum* L. cv. Laxton’s Progress 9 or Little Marvel) as described (Cline, 1986). Chloroplasts were suspended to 1 mg/ml chlorophyll in import buffer (IB, 50 mM HEPES-KOH, pH 8.0, 330 mM sorbitol) and kept on ice. Isolated thylakoids were obtained by osmotic lysis of intact chloroplasts. Briefly, suspensions of intact chloroplasts were centrifuged for 5 min at 1000 ×g, the supernatant discarded, and the chloroplast pellet was suspended at 1 mg/ml chlorophyll in lysis buffer (50 mM HEPES-KOH, pH 8.0, 10 mM MgCl_2_) and incubated on ice for 5 min. Lysates were then diluted with IB containing 10 mM MgCl_2_ followed by recovery of the thylakoids by pelleting at 3200 ×g for 8 min. Finally, thylakoids were resuspended in IB containing 10 mM MgCl_2_ for use in later assays.

### Import of *in vitro* translated radiolabeled cpTatC

cysteine-substituted cpTatC mRNAs were transcribed *in vitro* with SP6 polymerase (Promega) and were translated *in vitro* in the presence of ^3^H-leucine (Perkin Elmer) using a home-made wheat germ translation system. Translations were diluted 1:1 and adjusted with import buffer containing 30 mM leucine before use. For import, *in vitro* translated cpTatC was incubated with chloroplasts (0.33 mg chlorophyll/mL) and 5 mM Mg-ATP in import buffer with ~100 μE m^−2^ s^−1^ of white light in a 25°C water bath for 40 min. After import, intact chloroplasts were pelleted and resuspended in IB with 1 mM thermolysin for 40 min on ice. After proteolytic digestion, intact chloroplasts were re-purified by centrifugation through a 35% Percoll (GE Healthcare) solution in IB with 5 mM EDTA.

### Integration of *in vitro* translated unlabeled Hcf106 or Tha4

Cysteine-substituted Hcf106 or Tha4 mRNAs were transcribed *in vitro* with SP6 polymerase (Promega) and were translated *in vitro* in the presence of unlabeled leucine using a home-made wheat germ translation system. Translations were diluted 1:1 and adjusted to 30 mM leucine in IB. For integration assays, chloroplasts were first lysed, and thylakoids suspended in 10 mM MgCl_2_ IB. The *in vitro* translated Hcf106 or Tha4 were incubated with thylakoids (0.33 mg chlorophyll/ml) in IB with 10 mM MgCl_2_ in a 25 °C water bath for 20 min. After integration, thylakoids were washed with import buffer containing 10 mM MgCl_2_.

### Disulfide crosslinking between imported cpTatC and integrated Hcf106 or Tha4

1 mM copper phenanthroline (CuPhen) was added to catalyze disulfide formation between proximate cysteine residues. The 150 mM CuPhen stock solution contained 150 mM CuSO_4_ and 500 mM 1, 10-phenanthroline. The crosslinking reaction was carried out for 5 min before cessation by the addition of 50 mM N-ethylmaleimide (NEM; from a 1 M stock in ethanol) in 1×IB, 10 mM MgCl_2_. Thylakoids were recovered by centrifugation and were washed with 1×IB, 5 mM EDTA, and 10 mM NEM. Thylakoid pellets were re-suspended and divided into two halves for comparison of non-reducing and reducing SDS-PAGE. The non-reduced sample was resuspended in 100 mM Tris-HCl (pH 6.8), 8 M urea, 5% SDS, and 30% glycerol and the reduced portion was resuspended in 100 mM Tris-HCl (pH 6.8), 0.2 M DTT, 5% SDS, and 30% glycerol. Samples then were analyzed by SDS-PAGE and fluorography.

### Immunoprecipitation of crosslinked complexes

After crosslinking, thylakoid samples were dissolved with 1% SDS, 50 mM Tris-HCl, pH 7.6, 1 mM EDTA at 37 °C for 5 min and centrifuged for 5 min. Supernatant was transferred and diluted with 10 volumes of immunoprecipitation (IP) buffer (50 mM Tris-HCl, pH 7.6, 150 mM NaCl, 0.1 mM EDTA, 1% Triton X-100, and 0.5% deoxycholate). 30 μl of packed magnetic FLAG (α-DYKDDDDK) gel beads (Sigma) were added to the supernatant followed by end-over-end mixing at 4 °C, overnight. The beads were washed three times with IP buffer followed by a final wash of 0.05% Triton X-100 in Tris buffered saline (TBS). The bound protein was eluted with 0.1 M glycine HCl, pH 3.0, and then diluted with the same volume of sample buffer as eluate.

### Accession numbers

The sequence of the cpTatC template with the transit peptide and first 13 residues of the mature domain of the pea small subunit of ribulose-1,5-bisphosphate carboxylase/oxygenase can be found in NCBI GenBank under the accession no. AAA33685. The N-terminally truncated amino acid sequence (72-303) of cpTatC from *P. sativum* can be found in NCBI GenBank under the accession no. AAK93948. The mature sequence of Hcf106 from *P. sativum* can be found in NCBI GenBank under the accession no. AAK93949. The backbone structure of *A. aeolicus* TatC can be found in the RSCB PDB under the PDB code 4B4A.

## List of author contributions

Q.M. and C.D.S. conceived the study; Q.M. and C.P.N. collected and analyzed the crosslinking assay data; Q.M. wrote the article with contributions of all the authors; C.P.N. and C.D.S. edited and revised the original manuscript for publication; C.D.S. supervised and complemented the writing.

## Footnote

Carole Dabney-Smith is the author of contact and confirmed author listing and contributions to the work presented herein according to the COPE guidelines described in Plant Physiology Instructions for Authors.

## Acknowledgements

This work was supported by the Department of Energy: Office of Science Basic Energy Science (Award DE-SC0014441 to CDS). The authors wish to thank members of the Dabney-Smith lab for critical reading of the manuscript.

## Abbreviations

TAT: Twin Arginine Translocation
TM: transmembrane domain
APH: aminoacyl-tRNA synthetase
NEM: N-ethylmaleimide
CuPhen: Cu2+-1,10-phenanthroline
DTT: dithiothreitol
IB: import buffer

